# Escalating human exposure to tropical mosquito-borne viruses in Europe

**DOI:** 10.64898/2025.12.02.691392

**Authors:** Kyla Serres, Fabiana Gámbaro, Rosa Pietroiusti, Daniele Da Re, Maria F. Vincenti-Gonzalez, Darlan da Silva Candido, Elena Arsevska, Augustin Jacques de Dixmude, Dánnell Quesada-Chacón, Matthias Mengel, Dominik Paprotny, Raphaëlle Klitting, Cedric Marsboom, Wim Thiery, Guillaume Ghisbain, Diana Erazo, Simon Dellicour

## Abstract

Human-induced climate change has multiple public health impacts, including the expansion of the geographical range of vector-borne diseases. Pathogens such as dengue, chikungunya and Zika viruses, transmitted by *Aedes* mosquitos, can cause severe health outcomes ranging from acute febrile illness, chronic joint pain, to birth defects and even death. Evaluating the future risk of human population exposure is therefore crucial as large outbreaks could overwhelm healthcare systems. Europe, one of the fastest warming regions globally, harbours the competent mosquito vector *Aedes albopictus* in over 20 countries, making tropical *Aedes-*borne viruses an increasing threat to the continent, which has already experienced local outbreaks over the past two decades. Here we use an ecological niche modelling approach to assess past, present, and future risk of human population exposure to dengue, chikungunya, and Zika viruses in Europe. Our results show that recent climate change has already increased the potential exposure to these viruses, particularly across the Mediterranean basin, which is a current hotspot for local outbreaks. Major metropolitan areas in Spain, France, Italy, and Croatia are by now located in at-risk areas, and this risk is projected to intensify and expand northward by mid-century. Under a high greenhouse gas emissions scenario, European areas ecologically suitable for *Aedes*-borne virus circulation could increase by up to ∼70%, leading to an additional ∼50 million people living in areas at risk by the end of the century. These findings underscore the urgent need for strengthened vector and epidemiological surveillance, as well as preparedness strategies across newly suitable regions to anticipate future public health threats associated with these arboviral diseases.

## Introduction

Human activities, particularly fossil fuel use, are responsible for global warming^1^, with global temperatures already 1.2°C above pre-industrial levels as of 2024^2^. The ongoing human-induced climate crisis is driving severe and often irreversible impacts, including more frequent extreme weather events^3,4^, biodiversity and ecosystem degradation^5^, economic losses^6^, and profound effects on public health^7^. These public health impacts arise via multiple pathways, such as heat-related illness and injuries from extreme weather^8,9^, ecosystem-mediated effects, including increases in vector- and water-borne diseases^10,11^, socially-mediated effects (such as malnutrition and forced migration)^12^, and inequity-related pathways that disproportionately affect vulnerable populations^13^.

Vector-borne diseases are particularly sensitive to climatic conditions, as variations in temperature, precipitation and humidity directly influence the biological traits and distribution of the competent vectors^14–16^. Climatic conditions also shape the epidemiological and transmission dynamics of the pathogens they carry^17^, making vector-borne diseases among the most climate-responsive health threats globally^18^. Dengue (DENV), chikungunya (CHIKV), and Zika (ZIKV) viruses are transmitted to humans by mosquitoes of the *Aedes* (*Ae*.) genus, particularly *Ae. aegypti* and *Ae. albopictus* (also known as the “yellow fever mosquito” and the “Asian tiger mosquito”, respectively). While both species are expanding into temperate regions such as the United States, only *Ae. albopictus* is currently established and spreading across mainland Europe (with *Ae. aegypti* being currently restricted to Madeira and Cyprus), driven by human-mediated transports and increasingly favourable climatic conditions^19^.

European autochthonous outbreaks have become more frequent, particularly between May and November during the active mosquito season^20^. Since 2007, notable dengue and chikungunya virus outbreaks involving hundreds of locally-acquired cases have occurred in Croatia, Spain, Italy, and France^21^; and in 2019, the first autochthonous Zika virus outbreak was reported in Hyères, France^22^. The growing frequency of such outbreaks poses significant public health challenges. Dengue is often asymptomatic but can progress to severe disease forms, particularly in children, individuals with comorbidities^23^, and in cases of secondary heterotypic infections due to antibody-dependent enhancement (ADE), which can cause high fever, persistent vomiting, bleeding, and in some cases death^24^. Zika virus is also mostly asymptomatic; however, infection during pregnancy causes severe congenital malformations, such as microcephaly^25^. On the contrary, chikungunya infections are typically highly symptomatic, with debilitating joint pain that can persist for months or years^26^. Diagnosis is complicated due to the overall high proportion of asymptomatic infections as well as the overlapping clinical presentation with co-circulating arboviruses (such as the West Nile virus) and other febrile illnesses (such as COVID-19 and seasonal flu), which can result in missed or delayed treatment^27,28^. Large outbreaks may overwhelm healthcare systems^29^, while vector control efforts entail substantial economic and ecological costs^30^, further exacerbated by rising insecticide resistance in mosquito populations^31,32^.

Currently, no specific therapeutic (post-infection) treatment exists for these mosquito-borne diseases, and supportive care targeting symptom management remains the primary approach^33^. In the context of a warming European climate^34^, it is therefore essential to strengthen risk assessments for populations living in areas increasingly suitable for local circulation of vector-borne viruses across the continent, which can in turn help strengthen surveillance and mitigate future outbreaks. To address this, we train ecological niche models using autochthonous human infection records linked to local climate, land-use, and population variables to identify environmental covariates associated with local virus circulation, to highlight areas of high human exposure risk, and to evaluate the impact of climate and land-use changes on the evolution of human population exposure risk through time. Specifically, we use our ecological niche models to predict the spatio-temporal changes of areas at risk over the past century under both an observed historical climate and a counterfactual climate, the latter corresponding to a no climate change baseline that conserves the observed natural climate variability^35^. We then project the future evolution of the human exposure risk over the next century under three pathways that describe different greenhouse gas concentration trajectories alongside plausible socioeconomic development trends^36,37^. We find that climate change has already increased the risk of human exposure to *Aedes*-borne viruses in Europe, with future projections suggesting a northward expansion of ecologically suitable conditions for their local circulation. European areas ecologically suitable for the local circulation of Aedes-borne viruses could increase by up to ∼70% under a high emissions scenario and could even double under a very high emission scenario, underscoring the urgency of strengthening surveillance and preparedness measures across new ecologically suitable regions.

## Results

We model the risk of human population exposure to *Aedes*-borne viruses in Europe using records of autochthonous dengue, chikungunya, and Zika virus human infections reported at a subnational administrative level between 2007 and 2024. These occurrence records were associated with climate, land-use and population data (Table S1) to train ecological niche models using a boosted regression tree approach (see the Methods section for further detail). The resulting models estimate the ecological suitability at each location, i.e. to what extent the environmental condition in a given location could host a local circulation of *Aedes*-transmitted viruses leading to detected human cases. Hereafter simply referred to as “ecological suitability” for convenience, this metric ranging from 0 to 1 can be interpreted as a measure of the potential risk of human population exposure to these viruses.

To minimise the risk of overfitting, we employed a spatial cross-validation procedure and also evaluated several pseudo-absence (PA) sampling approaches with varying presence-to-PA ratios (Fig. S1, Table S2; see the Methods section for further detail). Given the heterogeneity in arbovirus surveillance across Europe, we ultimately opted for a surveillance-capability sampling approach^38^, which prioritises the sampling as pseudo-absences of administrative areas associated with relatively higher epidemiological surveillance scores and no confirmed case. Such a pseudo-absence sampling approach therefore allows taking into account spatial heterogeneity in epidemiological surveillance while minimising the risk of sampling pseudo-absences in areas with limited surveillance where infections could simply have gone undetected. Using this pseudo-absence sampling approach, we replicated our analyses and trained a total of 100 independent ecological niche models, which achieved good predictive performance; i.e. a mean area under the receiver operating characteristic curve (AUC) equal to 0.884 (95% confidence interval [CI] = [0.858, 0.876]) and prevalence-pseudoabsence-calibrated Sørensen’s indexes (SI_ppc_) values over 0.996 (95% CI = [0.995, 0.997]; Table S2).

To assess the relationship between each environmental factor and the estimated ecological suitability for a potential local circulation of *Aedes*-borne viruses, we evaluated how ecological suitability varies with individual predictors while holding others constant (Fig. S2). Furthermore, we also calculated and reported the relative influence of each environmental predictor in the trained ecological niche models. Climatic variables (air temperature, relative humidity, and precipitation) accounted for about 73% of the estimated ecological suitability, land-use for 21%, and human population for the remaining 6%. Estimated ecological suitability increased with higher fall temperatures (>12°C) and precipitation exceeding 3 mm/day, while human population counts were also positively associated with the estimated ecological suitability (Fig. S2). In contrast, high fall relative humidity (>80%) and areas with more than 10% of managed pasture cover were negatively associated with ecological suitability. Finally, the other predictors had negligible influence, as indicated by their associated relative influence estimates and plateaued response curves across their observed ranges (Fig. S2).

To evaluate the contribution of climate change to the establishment of *Aedes*-transmitted viruses in some European regions, we compared changes in ecological suitability simulated by ecological niche models forced with historical versus counterfactual climate data from the Inter-Sectoral Impact Model Intercomparison Project 3a (ISIMIP3a) reanalysis dataset 20Crv3-ERA5^35^. Compared to the historical data, the counterfactual data are based on the same environmental changes but a counterfactual climate where long-term trends have been removed. Across both the factual and counterfactual historical simulations, the Mediterranean basin consistently emerges as an area at risk (Fig. 1). However, under historical climate conditions, the simulations show marked increases in ecological suitability, with expansion along the Mediterranean, Adriatic, and Balkan coasts, as well as the Bay of Biscay (Fig. 1). While some regions, such as the Mediterranean basin, were already at risk, these results indicate that climate change has amplified the potential for human population exposure to *Aedes*-borne viruses to date. With most administrative areas being associated with a standard deviation below 0.2, our results were overall consistent across replicated analyses (Fig. S3).

**Figure 1.**
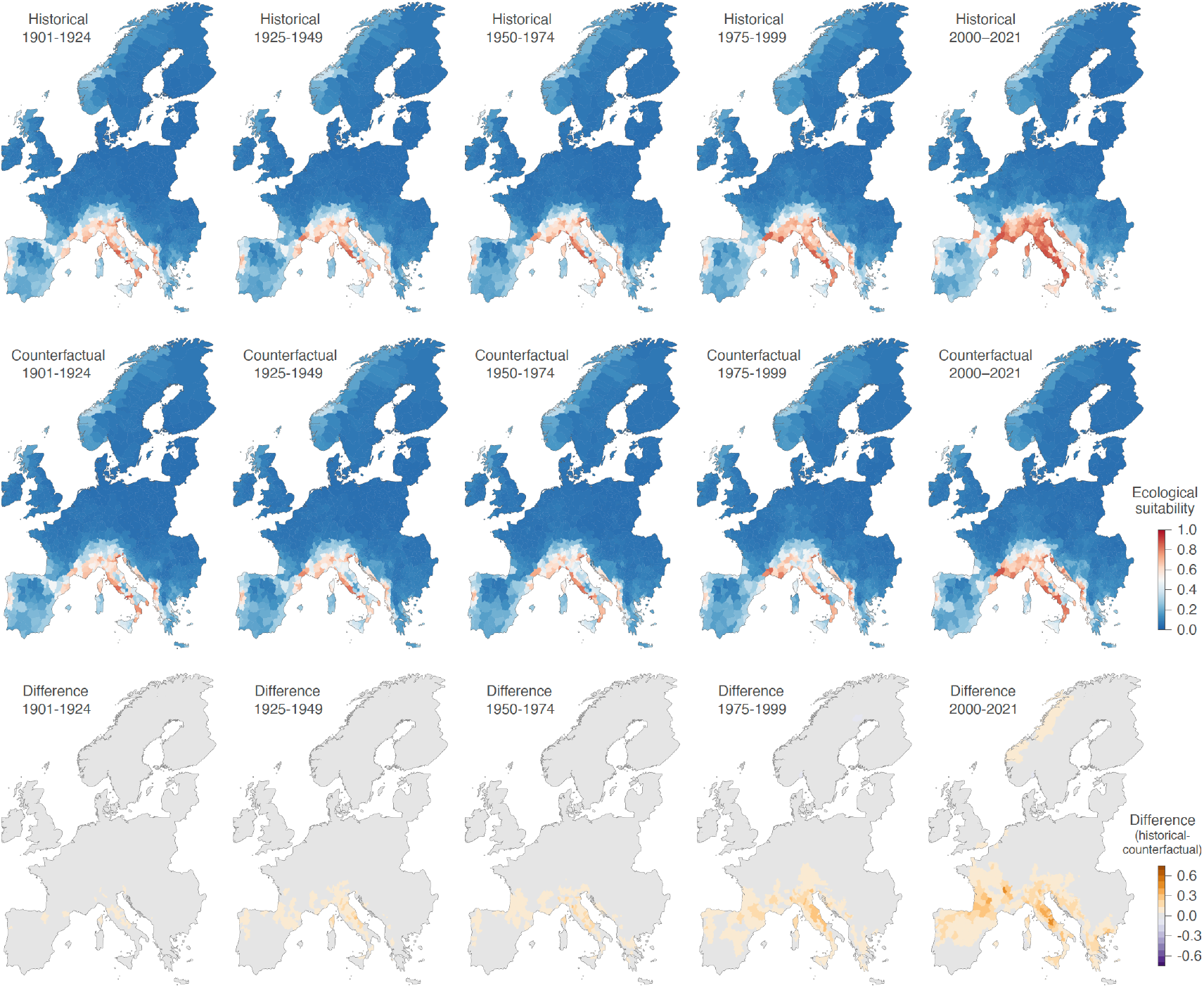
Changes in the areas ecologically suitable for the local circulation of *Aedes-*transmitted viruses (dengue, chikungunya, and Zika viruses) leading to detected human cases in Europe. Past ecological suitability is estimated for each administrative unit based on both a historical climate and a counterfactual baseline (i.e. a climatic scenario where long-term trends have been removed). Ecological suitability values are averaged over the estimates of 100 independent ecological niche models trained on present-day data retrieved from the ISIMIP3a reanalysis dataset 20CRv3-ERA5. The last row of maps displays the difference between the ecological suitability estimates obtained under the historical and counterfactual scenarios.

We then project how the risk of human population exposure to local circulation of *Aedes*-borne viruses may evolve in the next decades under different global change scenarios. For this purpose, we projected our trained ecological niche models across the 21^st^ century under three shared socioeconomic pathways (SSPs) corresponding to low, high, and very high greenhouse gas emissions^39,40^ (SSP1-2.6, SSP3-7.0, and SSP5-8.5; Figs. 2, S4). Currently, areas of high ecological suitability for the local circulation of *Aedes*-borne viruses are concentrated across southern Europe, particularly in Italy, the Mediterranean regions of southern France and northern Spain, the Adriatic coast and parts of the Bay of Biscay. Under the high (SSP3-7.0) and very high (SSP5-8.5) emissions scenarios, suitable ecological conditions expand northwards by mid-century (2050-2074), extending into central France, Belgium, and parts of the Netherlands. By the end of the century, this expansion could further intensify across the continent, areas ecologically suitable increasing by 68% (95% confidence interval = [58, 78]) under SSP3-7.0 and by 107% [89, 125] under SSP5-8.5 (Fig. 2). Northern countries such as Belgium and the Netherlands are projected to experience even larger increases, with areas ecologically suitable rising by 123% [103, 144] under SSP3-7.0. Several major populated cities, including Barcelona, Valencia, Paris, Marseille, Lyon, Rome, Milan, Naples, Ljubljana, Zagreb and Split, lie within these emerging or expanding high-risk zones. Toward the end of the century, the risk of human exposure to these viruses could further intensify across northern Europe under the very high climate change (SSP5-8.5) scenario, and some southern European regions, including parts of Italy, southern France, and northern Spain, show a decline in ecological suitability.

**Figure 2.**
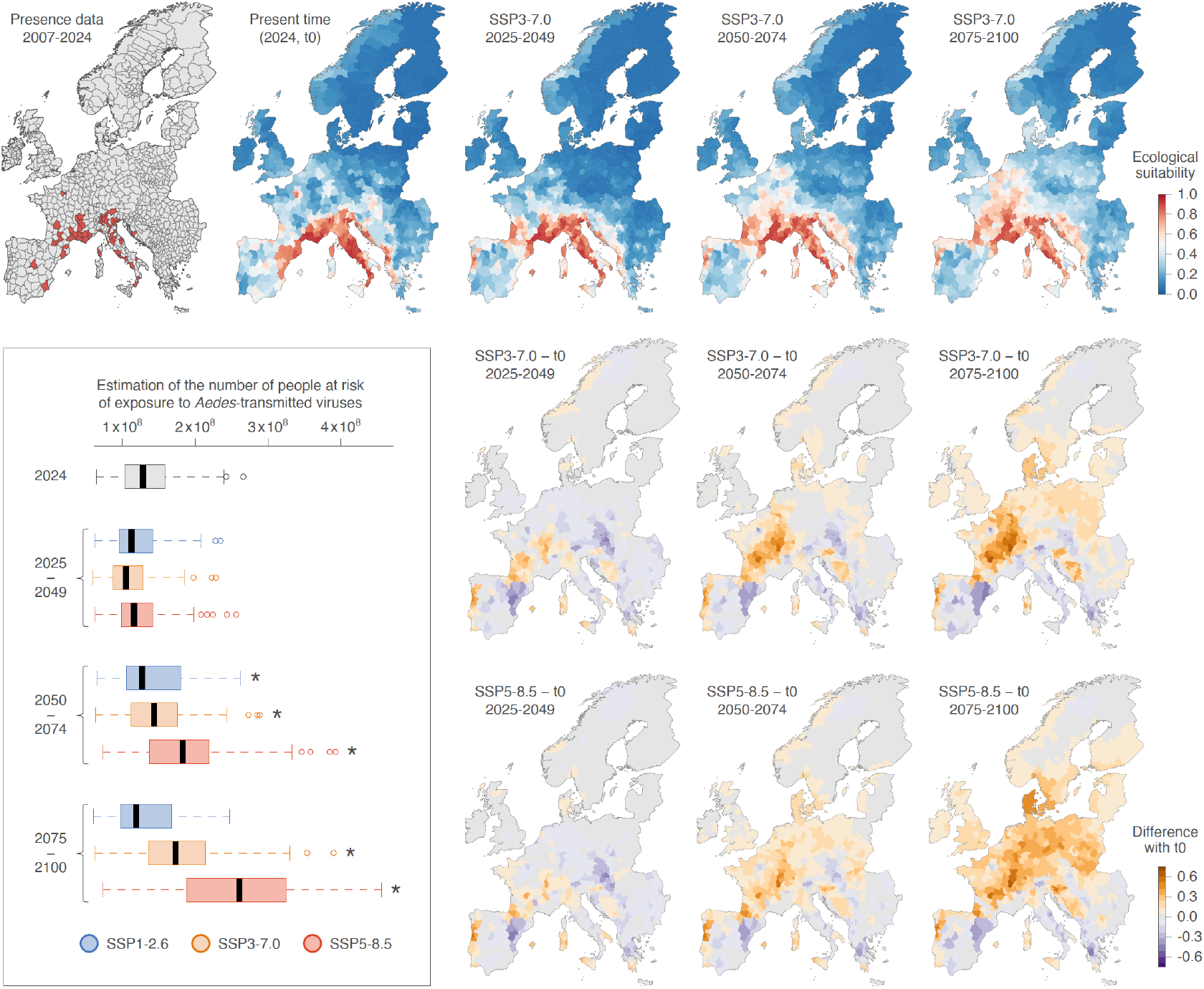
Future projections of the European areas ecologically suitable for the local circulation of *Aedes*-borne viruses and for the associated human population at risk of exposure under three different shared socio-economic pathways (SSPs). The first row of maps displays the spatial distribution of the presence data (administrative areas with confirmed local infection cases) from 2007 to 2024 used to train the ecological niche models, the predicted ecological suitability for the local circulation of those viruses at the present time (2024, also referred to as “t0”), and the future projections for three successive periods (2025-2049, 2050-2074, 2075-2100) under the high emissions scenario SSP3-7.0 (see Figure S4 for the projections under the low and very high emissions scenarios, SSP1-2.6 and SSP5-8.5). The second and third rows of maps display the difference in ecological suitability between estimates obtained for specific future projections for scenarios SSP3-7.0 or SSP5-8.5 and estimates obtained for the present-day (“t0”). Finally, the embedded boxplot graphic reports the number of people estimated to live in European areas at risk of exposure to local *Aedes*-borne virus circulation for the different periods and SSPs considered (see the Methods section for further detail on the approach used to define the areas at risk). (*) refers to a significant increase in the estimated number of people living in at-risk areas as compared to 2024 (as evaluated with a paired Wilcoxon test).

To quantify the evolution of the European population living in at-risk areas — defined as areas where ecological suitability estimates exceed the threshold value maximising the Sørensen’s Index (see Methods for further detail) — we combine ecological suitability projections with population counts projected under the corresponding global change scenario. For both mid- and late-century periods, the number of people living in at-risk European areas under SSP3-7.0 and SSP5-8.5 scenarios is significantly higher than in 2024 (paired Wilcoxon test; Fig. 2). Currently, approximately 137 million people [128, 146] reside in regions estimated as ecologically suitable for the local circulation of *Aedes*-borne viruses. Under a high emissions scenario (SSP3-7.0), the risk of human population exposure is projected to increase substantially, with an additional 56 million [49, 64] by the end of the century. Finally, under a very high emissions scenario (SSP5-8.5), the number of people living in at-risk areas is projected to nearly double by the end of the century, reaching a mean estimate of ∼260 million [243, 274] (Fig. 2). Because the increase in the population living in at-risk areas could also stem from the demographic growth itself, we also conducted these projections by fixing human population counts at their 2024 levels to isolate the contribution of the spatio-temporal evolution of the areas ecologically suitable for viral circulation. These alternative analyses yield to the same results, i.e. that the number of people living in at-risk areas in the second half of the century is projected to be significantly higher than in 2024 under both the SSP3-7.0 and SSP5-8.5 scenarios (paired Wilcoxon test).

## Discussion

Over the last two decades, several outbreaks of dengue, chikungunya, and Zika viruses have been reported across Europe, particularly in Spain, France, Italy, and Croatia^21^. These arboviruses pose growing public health challenges and currently lack specific antiviral treatments^33^. Vaccination options remain limited and context-dependent. For chikungunya, two vaccines exist, a live-attenuated vaccine (IXCHIQ)^41^, which was suspended in 2025 for people over 65^42^, and a virus-like particle vaccine (VIMKUNYA)^43^, which use is limited and primarily recommended for travellers at risk of exposure. For dengue, two vaccines are licensed: Dengvaxia, which is recommended only for individuals with prior dengue infection^44^ because of the risk of antibody-dependent enhancement and has been discontinued in several countries, and Qdenga, which can be administered regardless of previous infection and is now the major licensed vaccine in Europe^45,46^. In addition, a single-dose dengue vaccine – Butantan-DV – is currently in clinical trials and may provide immunological advantages over licensed vaccines through its use of more complete serotype-specific viral genomes^47^. Finally, no licensed vaccine is yet available for the Zika virus despite several candidates in clinical evaluation^48^.

Europe is the fastest-warming continent^49^, and climate-related risks like the expansion of tropical *Aedes*-borne viruses are increasingly threatening public health. Understanding both current and future suitability of the European environment for the circulation of these viruses is therefore critical to evaluate the evolving risk of human population exposure, which can in turn guide preparedness and mitigation strategies. In this study, we applied an ecological niche modelling approach to evaluate the contribution of climate change to the establishment of *Aedes*-borne viruses in Europe and to explore the evolution of the European population exposure risk under different global change scenarios. Our results indicate that climate change has likely been a contributor to the establishment of *Aedes*-borne viruses in several European regions. Our present-day estimations highlight southern European countries as potential hotspots for local circulation of *Aedes*-borne viruses, with modelled at-risk regions in France and Italy overlapping with areas affected by the chikungunya outbreak in the summer of 2025, which led to the detection of over 700 autochthonous cases in France and 350 in Italy^50,51^. Furthermore, our estimations for the risk of human exposure do not tend to significantly decrease even under the strong climate mitigation scenario (SSP1-2.6) — a pathway still technically possible but increasingly unlikely given current trajectories — suggesting that a substantial portion of climate-driven risk is already committed. Under a high emissions scenario (SSP3-7.0), ecologically suitable regions are projected to expand further into central and northern Europe and, under the very high emission scenario (SSP5-8.5), environmental conditions across Europe could become twice as suitable for the circulation of *Aedes*-borne viruses by the end of the century. In the latter scenario, southern regions may experience a decline in suitability, likely due to temperature increases and reduced water availability essential for mosquito development^52,53^. However, and although widely used for risk assessment, SSP5-8.5 is increasingly considered a less plausible scenario given existing climate policies, as it would entail warming approaching ∼5°C by 2100 and would require a reversal of current global mitigation efforts^54^.

Our study provides a comprehensive quantitative assessment focusing on the evolution of the human population exposure risk, and the results presented here are in line with several studies that previously aimed to assess *Aedes*-borne virus risk in Europe and globally. First, previous work using global suitability maps for *Aedes*-borne viruses has shown that combining occurrence records across multiple viruses improves risk prediction^38^. Other studies focusing on *Aedes albopictus* establishment and the timing of autochthonous dengue and chikungunya outbreaks across Europe found that increasingly favourable climatic conditions accelerate outbreak occurrence, with projections suggesting up to a five-fold increase in outbreaks under SSP5-8.5 by the 2060s^55^. Mechanistic and environmental-driven modelling has also demonstrated that *Aedes albopictus* expansion under climate change increases the potential for dengue transmission in western Europe^56^, with transmission expanding from the Mediterranean coasts into northern Spain and western France^56^. Global and regional risk mapping, as well as future projections, are increasingly recognised as essential tools for understanding the potential distribution and drivers of arboviral diseases and for informing public health planning^57^. In this context, our study evaluates the temporal evolution of the human exposure risk to *Aedes*-transmitted viruses, highlighting climate change as a factor that likely contributed to the establishment of those viruses in several southern European regions, particularly the Mediterranean basin, with further expansion expected toward northern Europe in the coming decades.

From a methodological perspective, we here explored different pseudo-absence sampling approaches, a critical consideration in ecological niche modelling because actual absence data are frequently unavailable or cannot be confidently inferred from under-surveyed areas. Given the spatial heterogeneity in arbovirus surveillance across Europe, we opted to sample pseudo-absences in areas where no human cases were reported but that are well-surveilled, which reduces the risk of sampling pseudo-absences in areas with undetected viral circulation. Specifically, administrative areas were sampled as pseudo-absence locations with a probability proportional to existing epidemiological surveillance score estimates^38^, which quantify the long-term probability that acute viral infections are detected and reported, accounting for differences in healthcare access, diagnostic capacity, and reporting practices.

Our future projections highlight regions that are becoming increasingly suitable for the circulation of *Aedes*-borne viruses, with potentially important public health implications across Europe. These findings underscore the urgency of strengthening integrated surveillance systems — both entomological and epidemiological — to enable early detection of local transmission and guide timely vector control responses. Because dengue infections can be asymptomatic in up to 80% of cases^58^, improved mosquito surveillance in current and potentially future ecologically suitable areas will be important for monitoring vector abundance and activity, although detecting *Aedes*-borne viruses directly in mosquitoes may remain challenging given the highly localised nature of transmission. Reliable and publicly accessible occurrence data from laboratories and hospitals are also critical to support robust modelling and risk assessment efforts.

As environmental suitability for the risk of local circulation of *Aedes*-borne viruses expands northward, diagnosis and clinical management may become more challenging, particularly in immunologically naïve populations. Continuous monitoring of both vector populations and viral circulation is therefore vital. Risk mapping analyses such as the ones presented in this study should also be regularly updated as new outbreak data emerge – especially from northern regions with climatic and land-use conditions distinct from southern areas – as these may reshape the projected risk patterns. Beyond surveillance, our results highlight the need for targeted and adaptive public health strategies. These could include citizen science initiatives for case detection and mosquito reporting, community education on mosquito prevention and individual vector control measures such as the use of repellents, window screens, and mosquito nets. With the potential for these viruses to become endemic in parts of Europe, additional safeguards — such as donor blood deferral, blood bank screening, and eventual vaccine deployment — may also need to be considered.

Finally, while the human exposure risk projections presented in this study provide an important foundation for anticipating future threats, further modelling efforts should integrate additional dimensions of transmission risk. These include human mobility from endemic regions, which remains the main pathway for viral introduction, and improved data on vector competence under European environmental conditions, which are still poorly understood. Integrating these elements will be crucial to refine model predictions and strengthen preparedness for *Aedes*-borne diseases in a rapidly changing climate.

## Methods

### Occurrence of human infection cases and environmental data acquisition

Occurrence data, i.e. confirmed and autochthonous (non-imported) cases of human infections with an *Aedes*-borne virus (DENV, CHIKV, or ZIKV), were collected and curated per NUTS3 (Nomenclature of Territorial Units for Statistics level 3) administrative area for a period of time ranging from 2007 to 2024 from multiple sources. These sources included occurrence records from the European Surveillance System (TESSy) database of the European Center for Disease Prevention and Control (ECDC); as well as peer-reviewed scientific articles^21^, national public health authorities (e.g., *Santé Publique France*), and the Platform for Automated extraction of Disease Information from the web (PADI-web), which is an automated biosurveillance tool that monitors online news sources to detect infectious disease events^59^. To account for variation in NUTS3 administrative unit sizes across European countries, we used optimised NUTS3 polygons developed by the European network for medical and veterinary entomology (VectorNet)^60^, which standardises unit sizes for spatial analyses. Given the sparsity of European occurrence data available for ZIKV and also, to some extent, for CHIKV, we opted for a joint ecological niche modelling analysis of all three *Aedes*-transmitted viruses considered here. The curated occurrence datasets were therefore merged to include all autochthonous occurrences of DENV, CHIKV and ZIKV infections for the subsequent ecological niche modelling analyses. In the context of sparse occurrence records, previous studies training ecological niche models for *Aedes*-borne viruses have demonstrated that combining occurrence records across multiple viruses could enhance predictive performance^38^.

Climate data (i.e. air temperature, precipitation, and relative humidity) were retrieved from the Inter-Sectoral Impact Model Intercomparison Project phase 3 (ISIMIP3)^61^. ISIMIP provides a standardised framework for climate impact simulations across multiple sectors, including health and vector-borne diseases. This framework allows for the assessment of how climate change affects systems such as health, water resources, and the economy by capturing the biophysical and socio-environmental processes underlying these impacts. ISIMIP3 is organised into two components: ISIMIP3a, which focuses on historical simulations driven by reanalysis-based observed climate and reconstructions of historical human forcings^35^, and ISIMIP3b which addresses future impact projections using climate forcings simulated by global climate models (GCMs), and projections of socioeconomic changes, under different scenarios^37^. Given the sensitivity of vector-borne diseases to seasonality, climate data were aggregated into seasonal means, which were defined as follows: winter (December of previous year, January, February), spring (March to May), summer (June to August), and autumn (September to November). Land-use data (i.e. croplands, pastures, urban areas, range lands, primary forested areas, secondary forested areas, primary non-forested areas and secondary non-forested areas) were obtained from the Land-Use Harmonization version 2 (LUH2) project^36^. Finally, human population data (i.e. population counts and population density) were obtained from the Compound Extremes Attribution of Climate Change (COMPASS) project^62^. Both land-use and population datasets were retrieved for past and future periods for scenarios matching those available in the ISIMIP framework, ensuring consistency with the climate simulations used in this study.

### Ecological niche modelling analyses and assessment of predictive performances

We estimated the risk of human population exposure to autochthonous *Aedes*-borne viruses by modelling the ecological suitability for a local circulation of those viruses that would lead to detected human cases. To this aim, we used a boosted regression tree (BRT) approach, which is based on a machine learning algorithm that generates an ensemble of sequentially fitted regression trees optimising the predictive probability of occurrence based on the environmental conditions at a specific location^63,64^. Such a predictive probability can here be interpreted as a measure of the risk of human exposure to *Aedes*-borne viruses, with values ranging from 0 (minimum risk) to 1 (maximum risk). We selected the BRT algorithm for its recognised ability to capture complex non-linear relationships between response and predictor variables, its robustness to outliers, and its superior predictive performance compared to other modelling techniques^63^.

Because epidemiological reporting intensity varies across countries and over time, the absolute number of reported cases cannot be used as a reliable proxy for disease prevalence. In this context, we used a Bernoulli BRT approach by considering a binary response variable made of presence and absence data. To this aim, administrative units with at least one confirmed autochthonous human case were treated as presence locations. Because the Bernoulli BRT approach requires both presence and absence data, we generated artificial absence data, commonly referred to as “pseudo-absences” (PA). Notably, in ecological niche modelling (ENM), one of the most critical factors influencing model performance is how well the presence and considered absence data reflect the actual relationships between the target organism and its environment^65^. As a result, PA sampling becomes a central focus in ENM because sampling PAs in areas where the target organism is actually present but not reported can logically impact the accuracy and predictive performances of the resulting ecological niche models. To reduce this risk, a variety of PA sampling procedures exist, ranging from simple random sampling to more sophisticated constraint-based approaches^66–68^. This topic remains widely debated within the ENM community, yet no universally accepted standard has emerged^69^. We here tested four different PA sampling approaches, as well as three different ratios between the number of administrative areas treated as presence locations and the number of PA ones: a 1:1 ratio (equal number of PA and presence locations), a 1:3 ratio (three times more PAs), and a 1:6 ratio (six times more PA). Additionally, to further reduce the likelihood of sampling PAs in areas where human infections occurred but remained undetected, we adopted a conservative approach by excluding all NUTS3 administrative units adjacent to an administrative unit treated as a presence location.

As introduced above, we tested four different PA sampling approach. The first one was a random sampling approach, in which PAs were randomly selected from the available pool of administrative units not associated with, nor adjacent to, known administrative units considered as presence locations. The second approach was an *Aedes albopictus* suitability weighted procedure where PAs were sampled according to the distribution of *Aedes albopictus* as predicted by Kraemer and colleagues^70^, giving higher selection probability to administrative areas with greater vector suitability. The third approach was a disease surveillance capability weighted procedure, in which PAs were sampled based on the predicted relative surveillance capability for emerging acute viral infectious diseases estimated by Lim and colleagues^38^. With this approach, administrative areas with higher surveillance capability scores were more likely to be sampled, accounting for spatial variation in disease detection and reporting. Finally, in the fourth approach, PAs were uniformly sampled within an environmental space using the method developed by Da Re and colleagues^68^ and implemented in the R package “USE”. This latest approach identifies the portion of the environmental space associated with suitable conditions from known presences using a kernel-based filter, and then uniformly samples pseudo-absences outside this region, thereby reducing the likelihood of introducing false absences into the training dataset. Figure S1 provides a visual example of each PA sampling approach for the different PA-to-presence ratios. For each PA sampling approach, we trained 30 independent BRT ecological niche models, each model using a different set of sampled PAs, ensuring variability in the training data. As detailed in the Results section, we eventually selected the third PA sampling approach based on relative surveillance capability, and for which we conducted 100 replicate BRT analyses. While all PA sampling approaches led to good predictive performances (Table S2), this PA sampling approach was eventually selected to minimise the risk of an impact of surveillance bias, such as the sampling pseudo-absences in areas with limited surveillance and undetected viral circulation. We also opted to consider an equal number of presences and PAs, consistent with evidence that it can improve performance in classification algorithms such as BRT^67^.

To train the ecological niche models, the temporal dimension of the occurrence data was explicitly accounted for: each presence record was linked to its reporting year, and the corresponding climate, land-use and population data (see Table S1 for all environmental factors included in our ecological niche model) for that year were assigned to the presence location. Infection records spanning multiple years were aggregated at each location and associated with the corresponding year range, with environmental data averaged over that period. If a gap of ten years or more occurred between presence records at the same location, the site was treated as two separate time points: one for the earlier time point, and one for the range after the gap. This approach avoids averaging across long periods without reported cases, which could potentially bias the resulting ecological niche models. Furthermore, each administrative area sampled as a PA was assigned a pseudo-year, sampled according to the probability distribution of the original presence years, and which was then used to assign to it the climatic predictor values corresponding to this pseudo-year.

We used the BRT algorithm implemented in the R package “dismo” version 1.3-16^71^. To address spatial autocorrelation and reduce the risk of model overfitting, we used a spatial rather than a standard cross-validation procedure, as the latter often overestimates model performance when occurrence data are spatially autocorrelated^64^. We employed a spatial cross-validation method based on the generation of spatial blocks, following the approach proposed by Valavi and colleagues as implemented in the R package “blockCV” version 3.1-5^72^. BRT models were trained using the following parameters: a tree complexity of 5, a learning rate of 0.001, and a step size of 10. We assessed model performance using two metrics: the area under the receiver operating characteristic curve (AUC) and the prevalence-pseudoabsence-calibrated Sørensen’s index (SI_ppc_). While the AUC is widely used, it has been criticised for its sensitivity to prevalence^10,73,74^, which motivated the inclusion of the SI_ppc_ as a complementary predictive performance metric. For each replicate BRT analysis, we computed the AUC and SI_ppc_; the latter following the threshold optimisation procedure outlined by Erazo and colleagues^10^ and which identifies the threshold that maximises the predictive performance (see also Ghisbain *et al*.^74^ for a similar approach). The AUC and SI_ppc_ metrics are all reported in Table S2 for the different PA sampling approaches and presence/PA ratios. We quantified the contribution of each environmental variable by computing their relative influence (RI) in the ecological niche models (as reported in Figure S2). For a given variable, the RI reflects how frequently it was used for tree splits, weighted by the squared improvement it contributed and averaged across all trees. We also generated response curves to illustrate how the ecological suitability estimates respond to changes in each environmental variable while holding all others at their median value (Fig. S2), providing insights into the model’s behaviour and predictor-response relationships.

### Evaluating the impact of climate change on the emergence of Aedes-borne viruses in Europe

We used climate forcing data from ISIMIP3a, which provides a standardised framework for assessing the impacts of historical climate variability and long-term trends across multiple sectors sensitive to climate and human management^35^. ISIMIP3a includes paired factual and counterfactual historical datasets: the factual dataset represents observed climate conditions derived from reanalysis products, while the counterfactual dataset represents a hypothetical climate without long-term changes (i.e., detrended climate, using the ATTRICI method^75^), preserving observed short-term variability. Comparing both datasets allows us to isolate the extent to which a phenomenon – in this case, the risk of human exposure to *Aedes*-borne viruses as quantified by the ecological suitability for the local circulation of those viruses leading to detected human cases – is driven by climate change. To assess the impact of climate change, we simulate the changes in past ecological suitability over the last century by using either factual or counterfactual climate data from the ISIMIP3a framework in our trained ecological niche models. These climate data were retrieved from the ISIMIP3a reanalysis dataset^76^ 20CRv3-ERA5, which combines ERA5 (1979-2024) with the Twentieth Century Reanalysis version 3 (20CRv3), homogenised to the ERA5 reference for the earlier period (1901-1978). Among the available ISIMIP3a reanalysis datasets, 20CRv3-ERA5 was selected as it provides the most up-to-date coverage of the 20^th^ and early 21^st^ centuries.

### Exploring the potential evolution of the risk of human exposure in a near future in Europe

Future climate projections were obtained from ISIMIP3b^77,78^, which provides bias-adjusted and harmonised outputs from global climate models (GCMs) participating in the Coupled Model Intercomparison Project Phase 6 (CMIP6)^37^. We used simulations from ten state-of-the-art GCMs – GFDL-ESM4, IPSL-CM6A-LR, MPI-ESM1-2-HR, MRI-ESM2-0, UKESM1-0-LL-2, CanESM5-2, CNRM-CM6-1, CNRM-ESM2-1, EC-Earth3, and MIROC6 – under three Shared Socioeconomic Pathways (SSPs) — SSP1-2.6, SSP3-7.0, and SSP5-8.5. These scenarios span a broad range of possible climate futures, from strong mitigation to high and very high emission trajectories. Within ISIMIP3b, the SSP5-8.5 scenario represents a worst-case warming pathway; SSP3-7.0 represents a world characterised by fragmented development and regional disparities; and SSP1-2.6 reflects a sustainability-oriented scenario consistent with ambitious mitigation efforts^37,79^. To estimate the future suitability of the European continent for the local circulation of *Aedes*-borne viruses, we projected the BRT models over the next decades under these three different SSP scenarios and ten GCMs. For each scenario and considered time period, estimated ecological suitability values at the administrative-area level were obtained by averaging projections across the 100 BRT repetitions for each GCM, and subsequently averaging these mean values across the 10 GCMs.

To quantify human exposure risk, we then combined projected ecological suitability with scenario-specific population data. For each scenario and period of time, we identified NUTS3 administrative units exceeding the model performance evaluation threshold (based on the SI_ppc_ threshold maximum) to classify areas as ecologically suitable or unsuitable. We then used population data of each climate scenario and summed the total population counts of all suitable administrative areas, yielding the number of people at risk. This procedure was repeated across 100 model replicates to capture the uncertainty associated with the model training step, and we eventually compared the distributions of the population at risk across scenarios and time periods to the present 2024 baseline using paired Wilcoxon tests.

## Code and data availability

R scripts related to the ecological niche modelling are all available at https://github.com/kserresbriggs/ATV_ENM_Europe.

## Acknowledgements

We thank the data providers of the European Center for Disease Prevention and Control (ECDC), the Platform for Automated extraction of Disease Information from the web (PADI-web), *Santé Publique France*, and the Inter-Sectoral Impact Model Intercomparison Project (ISMIP). We also thank Louise Chini for sharing the LUH2-GCB2024 land-use data.

## Author contributions

KS, FG, GG, DE, and SD: conceptualisation and methodology. KS, FG, and SD: formal analyses, validation. EA, KS, AJdD: epidemiological data collection and curation. RP, DQC, MM, and WT: climate data preparation. DP: population data preparation. KS, FG, SD: original draft writing and editing. RP, DRR, MFVG, DdSC, EA, DQC, MM, DP, RK, CM, and WT: manuscript editing and reviewing. GG, DE, and SD: study design, manuscript writing and editing.

## Ethics declarations

This study used aggregated, de-identified data and therefore did not require prior ethical approval. The authors declare no competing interests.

---

**Figure S1.**
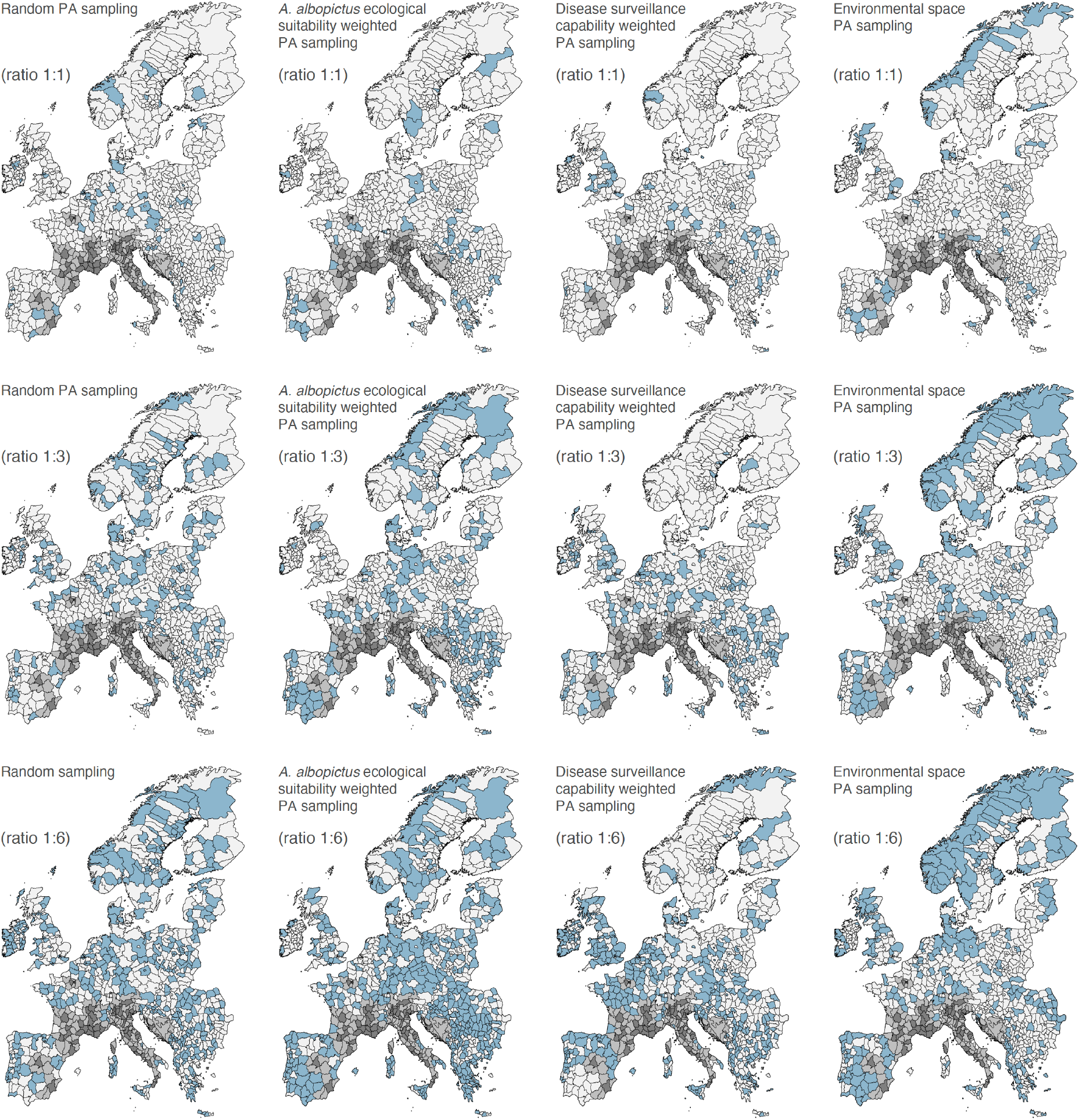
Visual comparison of the different pseudo-absences (PAs) sampling approach, as well as ratios between the number of presence and PA locations, considered for in ecological niche modelling analyses. For each PA sampling approach and considered presence-PA ratio, we here display one example of PA sampling corresponding to one boosted regression tree (BRT) analysis. On these maps, each administrative unit corresponds to a NUTS3 area (but see the Methods section for further detail on the optimised NUTS3 map). Light blue administrative units indicate sampled PAs, dark grey administrative units represent locations with a local virus circulation (presence units), and light grey administrative units denote units adjacent to presence units; both dark and light grey having been excluded from the set of possible PA areas (see the Methods section for further detail).

**Figure S2.**
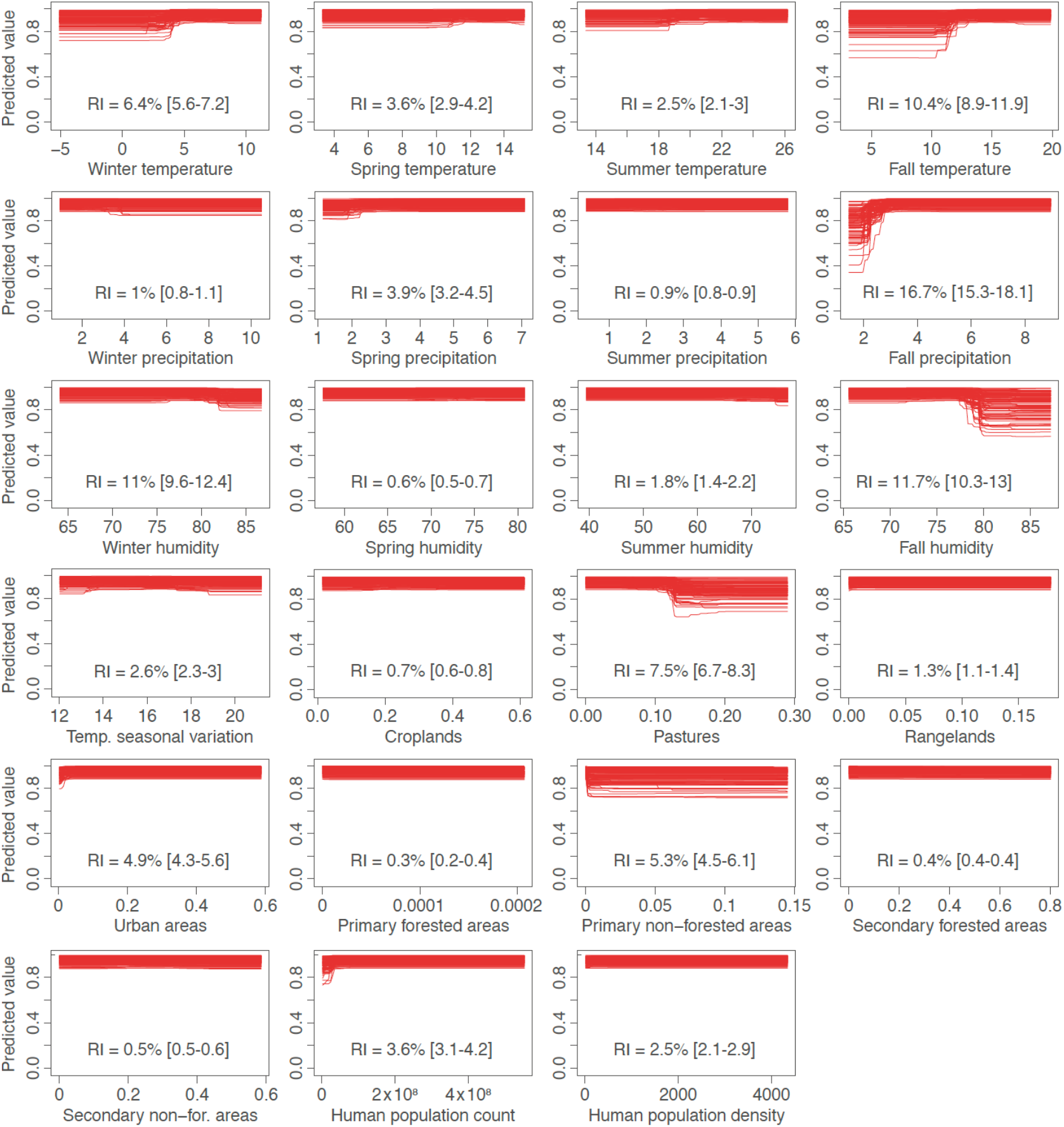
Response curves and relative influences (RI) of the ecological niche models. For each environmental predictor, we report the response curves obtained from 100 replicate boosted regression tree (BRT) analyses. These response curves depict the estimated relationship between each predictor and the ecological suitability for the local circulation of *Aedes-borne* viruses.

**Figure S3.**
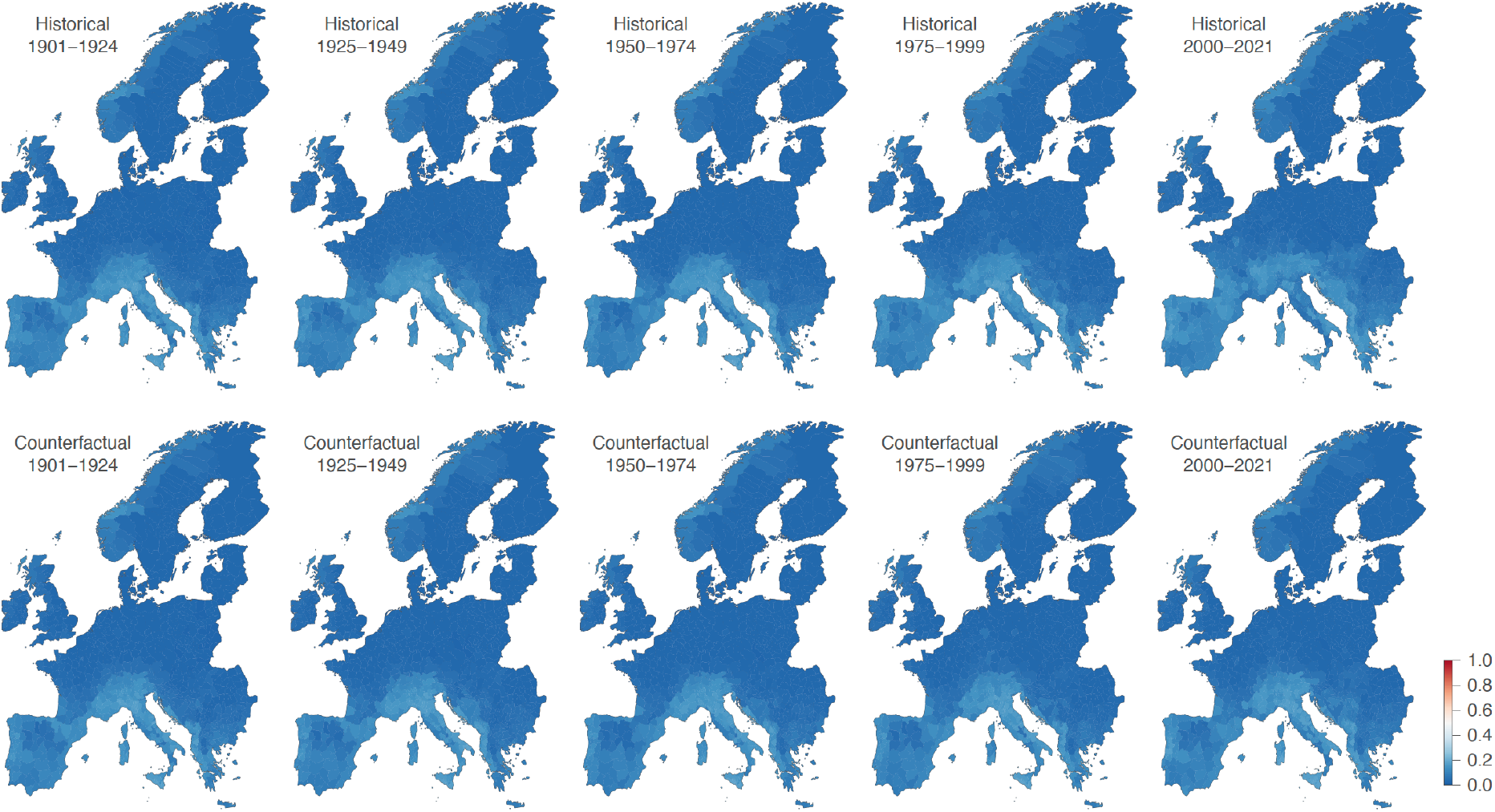
Uncertainty associated with the ecological suitability estimations reported in Figure 1. Specifically, we here report the standard deviation computed, for each administrative unit, from the 100 independent boosted regression trees (BRT) analyses.

**Figure S4.**
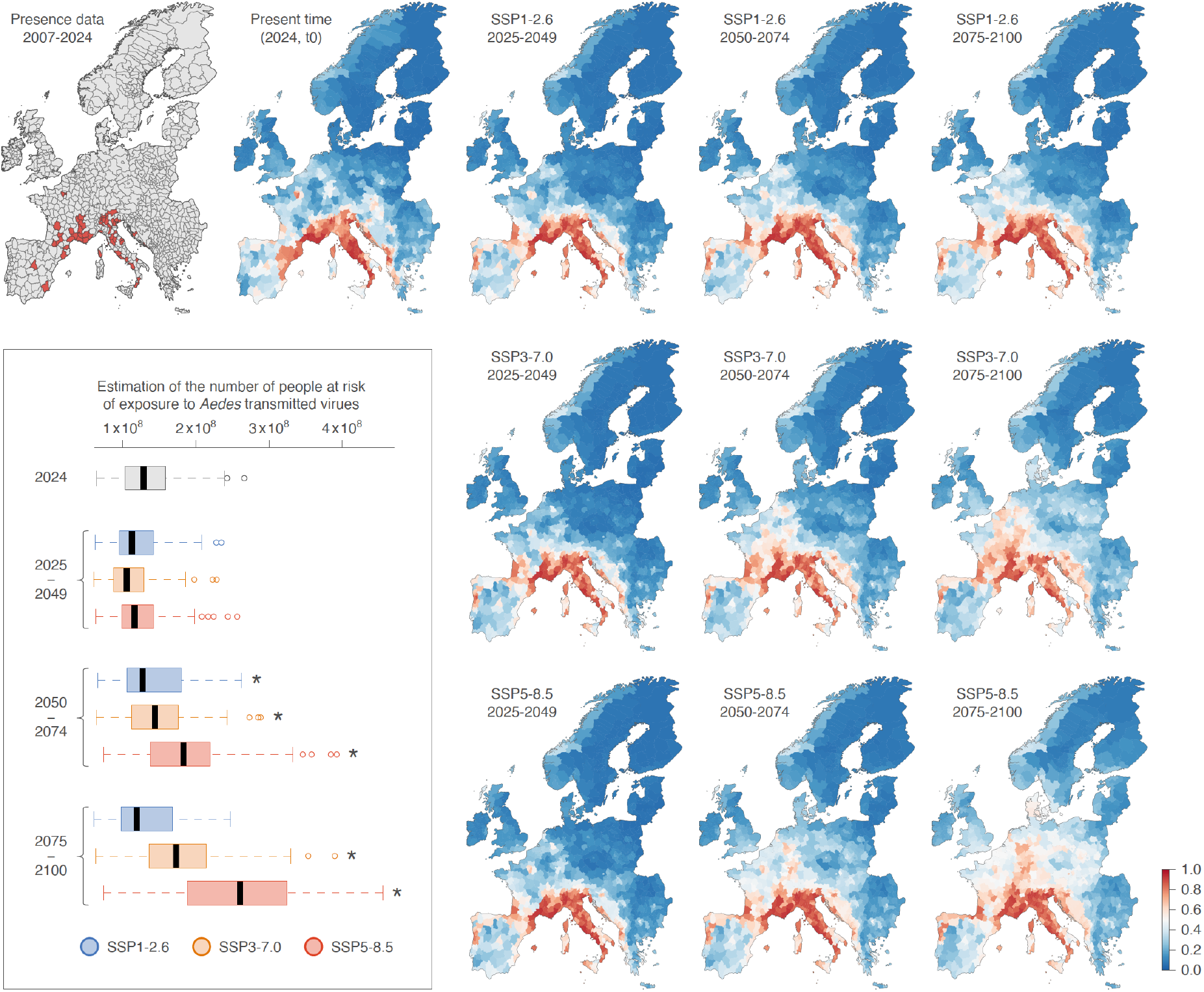
Future projections of the European areas ecologically suitable for the local circulation of *Aedes*-borne viruses and for the associated human population at risk of exposure under three different sShared socio-economic pathways (SSPs). This figure is an alternative version of Figure 2 solely focusing on the absolute ecological suitability values estimated for the different SSPs and future periods of time. The first row of maps displays the spatial distribution of the presence data (administrative areas with confirmed local infection cases) from 2007 to 2024 used to train the ecological niche models, the predicted ecological suitability for the local circulation of those viruses at the present time (2024, also referred to as “t0”), and the future projections for three successive periods (2025-2049, 2050-2074, 2075-2100) under the low emission scenario SSP1-2.6. The second and third rows of maps display the future projections for three successive periods (2025-2049, 2050-2074, 2075-2100) under the high and very high emission scenario, SSP3-7.0 and SSP5-8.5, respectively. As in Figure 2, the embedded boxplot graphic reports the number of people estimated to live in European areas at risk of exposure to local *Aedes*-borne virus circulation for the different periods and SSPs considered (see the Methods section for further detail on the approach used to define the areas at risk).

**Table S1.** Environmental factors included in the ecological niche modelling analyses. “ISIMIP” and “LUH” refer to the Inter-Sectoral Impact Model Intercomparison Project and Land-Use Harmonization databases, respectively.

| Environmental factors | Unit | Spatial resolution | Temporal resolution | Source | URL |
| --- | --- | --- | --- | --- | --- |
| Air temperature | °C | 0.5°x0.5° | monthly mean | ISIMIP3 | https://www.isimip.org/protocol/3/ |
| Temperature seasonal variation | °C | 0.5°x0.5° | monthly mean | ISIMIP3 | https://www.isimip.org/protocol/3/ |
| Precipitation | mm/day | 0.5°x0.5° | monthly mean | ISIMIP3 | https://www.isimip.org/protocol/3/ |
| Near surface relative humidity | % | 0.5°x0.5° | monthly mean | ISIMIP3 | https://www.isimip.org/protocol/3/ |
| Croplands | % | 0.5°x0.5° | annual | LUH2 | https://luh.umd.edu/ |
| Pastures | % | 0.5°x0.5° | annual | LUH2 | https://luh.umd.edu/ |
| Urban areas | % | 0.5°x0.5° | annual | LUH2 | https://luh.umd.edu/ |
| Range lands | % | 0.5°x0.5° | annual | LUH2 | https://luh.umd.edu/ |
| Primary forested areas | % | 0.5°x0.5° | annual | LUH2 | https://luh.umd.edu/ |
| Secondary forested areas | % | 0.5°x0.5° | annual | LUH2 | https://luh.umd.edu/ |
| Primary non forested areas | % | 0.5°x0.5° | annual | LUH2 | https://luh.umd.edu/ |
| Secondary non forested areas | % | 0.5°x0.5° | annual | LUH2 | https://luh.umd.edu/ |
| Human population | population counts | 0.5°x0.5° | annual | COMPASS | https://compass-climate.eu/ |
| Human population density | people/km | 0.5°x0.5° | annual | COMPASS | https://compass-climate.eu/ |

**Table S2.** Predictive performance of ecological niche models trained with different pseudo-absence (PA) sampling approaches and ratios between the number of presence and sampled PA locations. Predictive performance metrics include the area under the curve (AUC) and the prevalence-pseudo-absence calibrated Sørensen index (SI_ppc_). For each PA sampling approach and ratio, 30 independent replicate boosted regression tree (BRT) models were trained; and we here report the mean value of these two predictive performance metrics computed among replicates, as well as the associated 95% confidence intervals (CIs). (*) As detailed in the text, we eventually opted for the PA sampling approach based on disease surveillance capability as well as for a presence/PA ratio of 1:1, for which the 100 replicated BRT analyses yielded to a mean AUC of 0.867 (95% CI = [0.858, 0.876]) as well as a mean SI_ppc_ of 0.996 (95% CI = [0.995, 0.997]) associated with a mean threshold value maximising the SI_ppc_ equal to 0.462 (95% CI = [0.443, 0.481]).

| Pseudo-absence (PA) sampling approach | Presence/PA ratio | Area under the curve (AUC) | Prevalence-pseudo-absence calibrated Sørensen index ($SI_{ppc}$) | Threshold value maximising the $SI_{ppc}$ |
| --- | --- | --- | --- | --- |
| Random PA sampling | 1:1 | 0.885 [0.872, 0.897] | 0.997 [0.995, 1.000] | 0.421 [0.383, 0.460] |
|  | 1:3 | 0.883 [0.871, 0.895] | 0.982 [0.977, 0.987] | 0.387 [0.372, 0.403] |
|  | 1:6 | 0.884 [0.872, 0.896] | 0.971 [0.965, 0.977] | 0.292 [0.277, 0.308] |
| *Aedes albopictus* suitability weighted PA sampling | 1:1 | 0.879 [0.862, 0.895] | 0.996 [0.993, 0.999] | 0.421 [0.387, 0.455] |
|  | 1:3 | 0.861 [0.846, 0.875] | 0.973 [0.966, 0.981] | 0.340 [0.315, 0.365] |
|  | 1:6 | 0.877 [0.865, 0.889] | 0.950 [0.943, 0.958] | 0.293 [0.275, 0.311] |
| Disease surveillance capability PA sampling | 1:1* | 0.884 [0.864, 0.904] | 0.998 [0.996, 1.000] | 0.418 [0.387, 0.449] |
|  | 1:3 | 0.872 [0.860, 0.884] | 0.978 [0.973, 0.984] | 0.362 [0.343, 0.381] |
|  | 1:6 | 0.876 [0.864, 0.887] | 0.958 [0.953, 0.964] | 0.281 [0.266, 0.295] |
| Environmental space PA sampling | 1:1 | 0.848 [0.829, 0.867] | 0.997 [0.995, 1.000] | 0.451 [0.422, 0.480] |
|  | 1:3 | 0.861 [0.844, 0.877] | 0.983 [0.979, 0.987] | 0.376 [0.358, 0.394] |
|  | 1:6 | 0.879 [0.863, 0.895] | 0.970 [0.965, 0.976] | 0.330 [0.307, 0.354] |

